# Intestinal challenges shape the polarisation of protective dural memory CD4 T cells

**DOI:** 10.1101/2025.09.09.673536

**Authors:** Aaron Fleming, Rafael Di Marco-Barros, David A. Posner, Colin Y.C. Lee, Andrew Stewart, Zewen Kelvin Tuong, Karen Neish, Ana Peñalver, Mia Cabantous, Nathan Richoz, Anais Portet, Katherine Harcourt, Eleanor Gillman, David Ruano-Gallego, Gad Frankel, David Withers, Simon Clare, Menna R. Clatworthy

## Abstract

The meninges house several innate and adaptive immune cell populations^1–3^. These predominantly localise within the dura mater and include gut-derived IgA-secreting plasma cells^4^. Whether T cell adaptive memory in the dura is similarly linked to the gut is currently unknown. Here we show that dural CD4 T cell polarisation to a T helper (Th) 1, Th2 and Th17 state is determined by the nature of the immunological challenge encountered in the gastrointestinal tract. We find that intestinally polarised CD4 T cells seed to the dura in a CXCR6-CXCL16-dependent manner, express tissue-residency markers and are long-lived. Functionally, these orally-primed dural CD4 T are capable of a rapid antigen-specific recall response that limits pathogen spread into the brain following intravenous re-challenge. Our work reveals how linked intestinal and dural immunity enables the central nervous system to accrue immunological memory of gut microbes, the most likely source of life-threatening bloodborne pathogens.

## Introduction

The dura mater, the outer meningeal layer, contains a vascular network that includes fenestrated endothelium, presenting a route by which circulating microbes can access the central nervous system (CNS)^5–7^. Multiple populations of immune cells congregate around the dural venous sinuses^1–4^, whilst specific macrophage populations lie in close apposition to the meningeal vasculature^8^. In additional to potential roles in defending the CNS from infectious challenges, meningeal immune cells also have the capacity to influence brain function via cytokine production^9–12^.

In the periphery, including within the intestine, CD4 T cell polarisation to differing T helper cell states occurs following activation and the provision of signals that define the immunological context, including antigen presenting cell-derived cytokines^13^, microbial metabolites^14,15^ and strength of TCR signal^16^. This generates TBET^+^ IFNγ-producing Th1 cells, GATA-3^+^ IL4/13-producing Th2 cells or RORγt^+^ IL-17/22-producing Th17 cells, tailored to the prevalent immunological challenge^17^. Whether dural CD4 T cell polarisation can be influenced by systemic, or even gastrointestinal infection is currently unknown, but an important question given the proven effects of T cell-derived cytokines on brain function.

### Intestinal challenge shapes the polarisation of dural CD4 T cells

Given that the key cellular effectors of dural humoral memory, antibody-secreting plasma cells, may be educated in the gut, we hypothesised that gut-activated and polarised CD4 T cells, the orchestrators of cellular adaptive immunity, may similarly re-locate to the dura. In steady state, αβ CD4 T cells formed a minority population among dural immune cells (**Fig. S1A**) and were predominantly non-naïve, with around half of cells expressing TBET (**Fig. 1A, S1 B-E**). Following oral challenge with dextran sodium sulphate (DSS) (**Fig. 1B**), an osmotic colitogen that induces epithelial barrier breach and expansion of intestinal Th17 cells^18^, we observed a marked increase in RORγt^+^ Th17 cells in the colon (as expected), but also within the extravascular compartment of dura, in contrast to circulating CD4 T cells and those in the spleen (**Fig. 1C-F, S2 A-B**). Similarly, 21 days after oral challenge with *C. rodentium*, an attaching-effacing pathogen known to induce the expansion of intestinal Th17 cells^19^, we observed a significant increase in colonic and extravascular dural RORγt-expressing CD4 T cells (**Fig. 1G-H, S2 C-F**). These dural Th17 cells were predominantly IL-17-producing, with little intracellular IL-22 identified (**Fig. 1I-J, S2 I**). This increase in IL-17a-expressing cells was specific to CD4 T cells, and was not observed in the γδ or CD8 T cell compartments within the dura (**Fig. S2 G-H**). Spatially, Th17 cells were located along the dural venous sinus (**Fig. 1K**).

**Figure 1.**
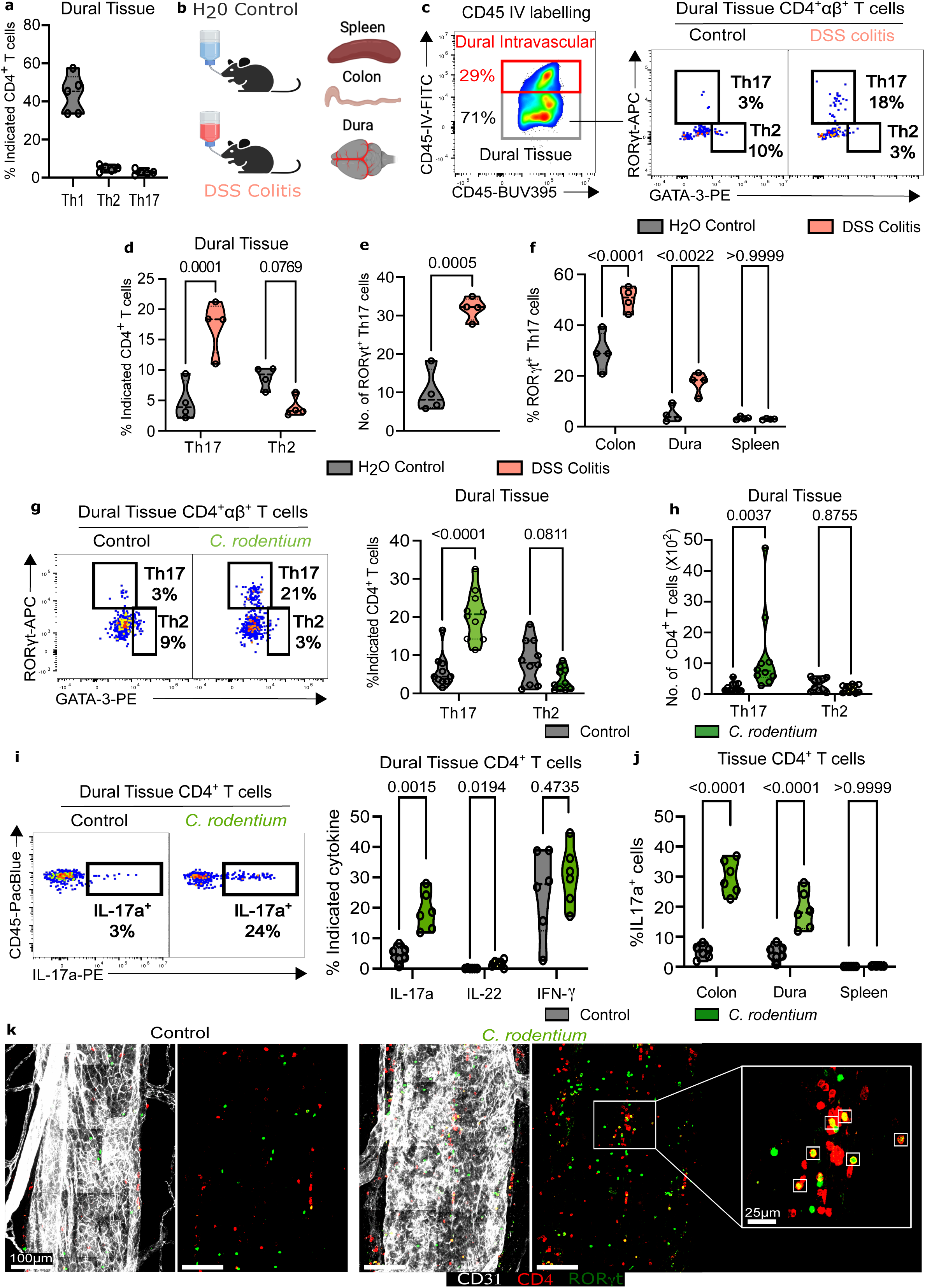
Dural T cells can be polarised by intestinal challenges to a Th17 phenotype. **A**: Th cell subset representation in extravascular (CD45IV^neg^) dural tissue. **B**: Schematic of DSS colitis model. Mice received 2.5% dextran sodium sulfate (DSS) in drinking water for two periods of 5 days separated by a 14-day interval and were culled at day 31. **C**: (left) Representative gating strategy used to discriminate dural intravascular (CD45-IV^+^) versus tissue (CD45-IV^-^) immune cells and (right) representative cytometry of dural Th cell subsets (*n* = 4). **D**: Cumulative data of flow in (c) (*n* = 4). **E**: Bead-normalised quantification of total CD4^+^αβ^+^Rorγt^+^ Th17 cells from dural tissue of control versus DSS colitis mice *(n* = 4). *P* value calculated using two-tailed Student’s unpaired t-test **F**: Representation of CD4^+^αβ^+^RORγt^+^ Th17 cells amongst CD4^+^ T cells from indicated organs (*n* = 4 per group). **G**: Representative cytometry (left) or cumulative violin plots (right) showing the representation of Th17 and Th2 cells amongst total CD4^+^ T cells from dural tissue of control versus *C. rodentium*-infected mice 21 dpi (*n* = 10). **H**: Bead-normalised quantification of total CD4^+^αβ^+^Rorγt^+^ Th17 and CD4^+^αβ^+^Gata-3^+^ Th2 cells from dural tissue of control versus *C. rodentium*-infected mice 21 dpi (*n* = 10). **I**: Representative cytometry (left) and cumulative violin plots (right) showing percentage of dural CD4^+^ T cells from *C. rodentium*-infected versus control mice expressing the indicated cytokine upon stimulation *ex vivo* (*n* = 6). *P* values calculated using multiple unpaired student *t*-test with Bonferroni-Dunn multiple comparisons correction. **J**: Percentage of CD4^+^ T cells in the indicated organs isolated from *C. rodentium*-infected versus control mice expressing IL-17 upon stimulation *ex vivo* (*n* = 6). **K**: Whole mount confocal microscopy of the superior sagittal sinus from control (left) and *C. rodentium* (right)-infected mice. Th17 cells highlighted in white squares. *P* values were derived using two-way ANOVA with Bonferroni’s post-test unless otherwise stated.

To determine if the linked expansion of gut and dural CD4 T cells was limited to the context of a Th17-polarising intestinal stimulus, we next challenged mice with the helminth *Schistosoma mansoni*^20^. *S. mansoni* is a worm with a complex life-cycle, initially infecting via cercariae, which burrow into the skin, transform into schistosomula, and then enter the vasculature and migrate to the portal system, where they mature into adult worms. Eggs released by female parasites within the vasculature extravasate, traverse the intestinal wall, and ultimately cross the epithelium into the lumen of the intestine where they induce a strong Th2 response^21^. At 42 days following challenge with *S. mansoni*, we found a marked expansion of GATA-3-expressing Th2 cells in the intestine, with a similar expansion observed in the dura, but little change in splenic Th2 cells (**Fig. 2A-B, S3 A-B**). Confocal imaging of dural wholemounts confirmed a substantial increase in CD4 T cells, particularly GATA-3-expressing cells, along the walls of the venous sinuses (**Fig. 2C**). In contrast, following oral challenge and the establishment of gut-limited chronic infection with *Trichuris muris* (mouse whipworm, known to induce a dominant Th1 response in C57BL/6 mice^22^), an increase in TBET-expressing Th1 cells number was present in the colon and dura (**Fig. 2D-E, S3 C-D**).

**Figure 2.**
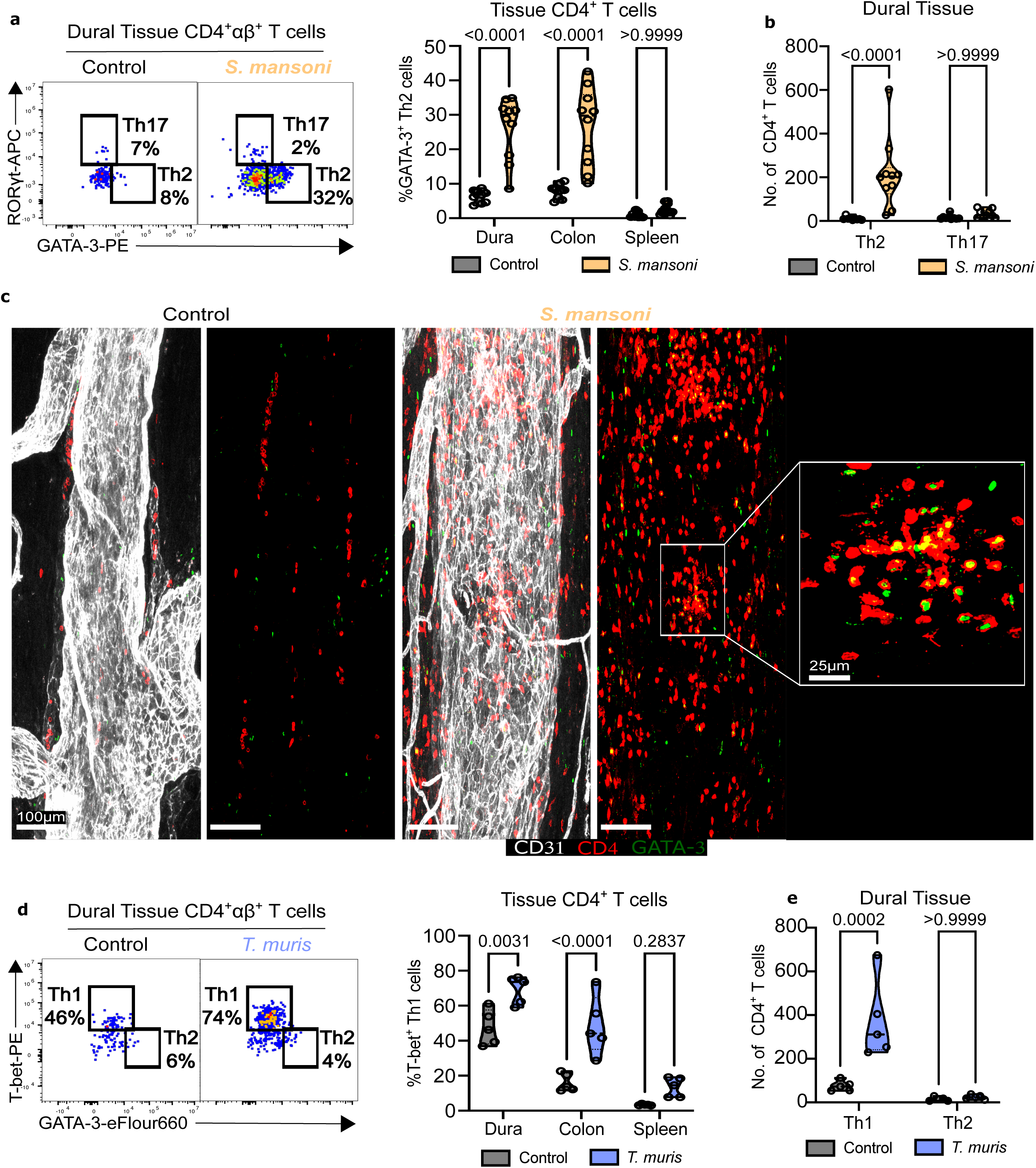
Dural T cells can be polarised by intestinal challenge to a Th2 or Th1 phenotype **A**: Representative cytometry (left) and cumulative violin plots (right) showing the representation of Th17 and Th2 cells amongst total CD4^+^ T cells from the indicated tissue of control versus *S. mansoni*-infected mice 42 dpi (*n* = 9-10). **B**: Bead-normalised quantification of total dural Th17 and Th2 cells from control versus *S. mansoni*-infected mice 42 dpi (*n* = 9-10). **C**: Whole mount confocal microscopy of the superior sagittal sinus from control (left) and *S. mansoni* (right)-infected mice 42 dpi. **D**: Representative cytometry (left) and cumulative violin plots (right) showing the representation of Th1 and Th2 cells amongst total CD4^+^ T cells from the indicated tissue of control versus *T. muris* infected mice 21 dpi (*n* = 5). **E**: Bead-normalised quantification of total dural Th1 and Th2 cells from control versus *T. muris* infected mice 21 dpi (*n* = 5). *P* values were derived using two-way ANOVA with Bonferonni’s post-test unless otherwise stated

Altogether our data show that the nature of an orally administered intestinal infection not only determines gut T cell polarisation, but can also profoundly alter the polarisation of tissue, extra-vascular dural αβ CD4 T cells.

### Intestinal challenge leads to expansion of antigen-specific CD4 T cells in the dura

To further probe the relationship of T cells in the gut and dura, we performed T cell receptor (TCR) sequencing on paired samples of dura, duodenum, ileum, proximal and distal colon 21 days following the induction of DSS colitis (**Fig. 3A**). We found a significant increase in the proportion of identical TCRs shared between dura and duodenum and distal colon in DSS mice compared with water-treated controls (**Fig. 3B-C**), suggesting that clones expanded in the intestine may seed to the dura. To test this further, we orally challenged mice with an attenuated 2W1S peptide-expressing *Salmonella enterica* serovar typhimurium (*S.* typhimurium), which is intestinal-limited during early infection^23^. At 21 days following infection, 2W1S-specific, CD62L-negative CD4 T cells were expanded in the extravascular compartment of both colon and dura, and to a much lesser extent in the spleen (**Fig. 3D-E**). These dural CD4 T cells were CD44^+^, expressed the tissue-residency marker CD69 (**Fig. S4 A-C**), and were IFNγ-secreting (**Fig. 3 F-G).** Together, these data support the conclusion that intestinal polarised, antigen-specific CD4 T cells relocate to the dura to establish a tissue memory population at this CNS border tissue following gut challenge.

**Figure 3.**
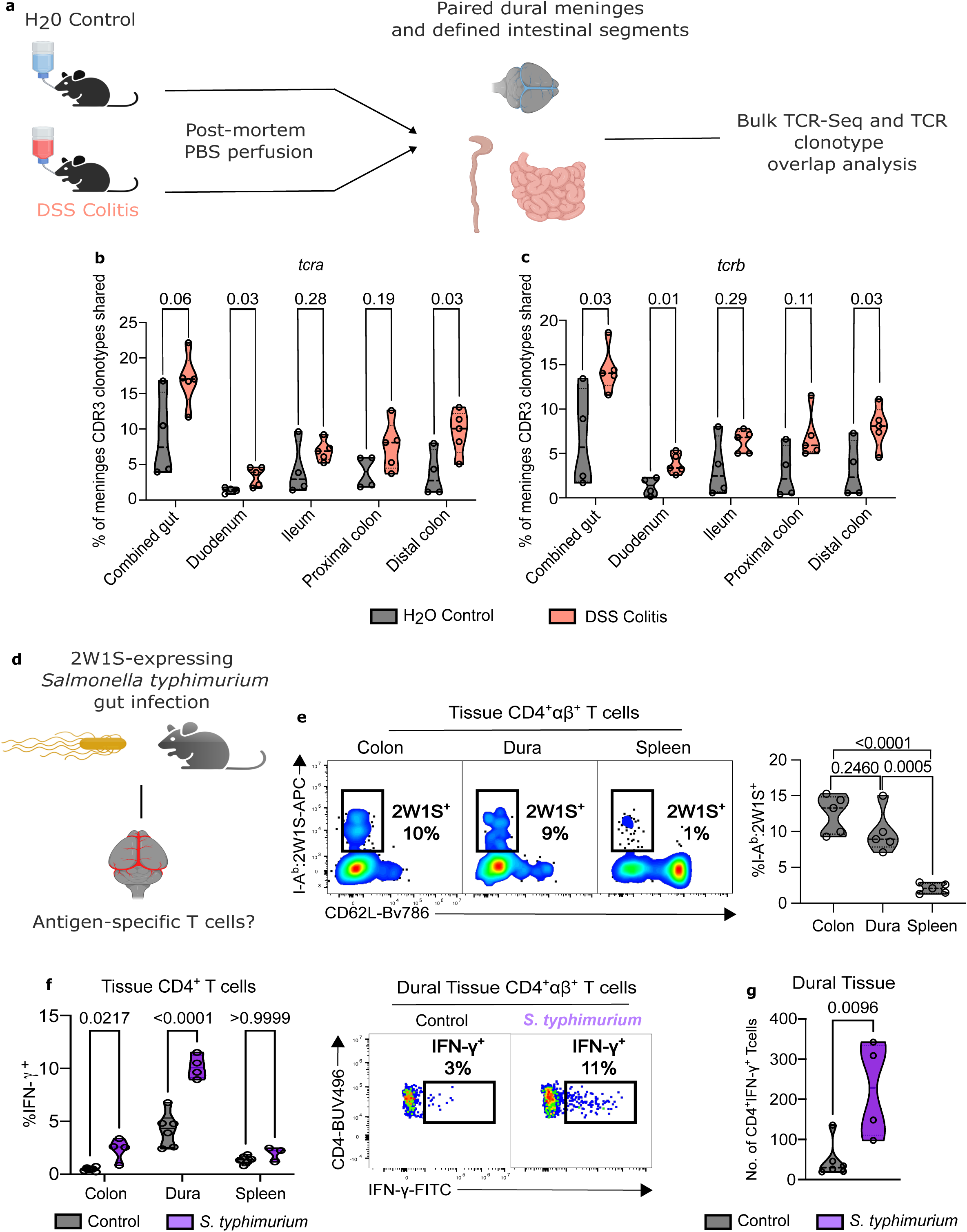
Intestinal challenge leads to accumulation of antigen-specific CD4 T cells in the dura **A**: Experimental design for TCR clonotype analysis in DSS colitis. Mice received 2.5% DSS or control H_2_O in their drinking water for 5 days and were sacrificed 16 days later with PBS perfusion. Dural meninges with paired intestinal segments were harvested for TCR-seq analysis. **B, C**: Percentage of dural *tcra* (b) or *tcrb* (c) CDR3 clonotypes shared with the indicated intestinal segments (*n* = 4-5 per group). *P* values calculated using multiple Mann-Whitney U tests with Bonferroni-Dunn multiple comparisons correction. **D**: Experimental set-up to identify dural antigen-specific CD4^+^ T cells in response to oral infection with *S.* typhimurium-2W1S. Mice were intravenously labelled with anti-CD45 and culled 21dpi. **E**: Representative cytometry (left) and cumulative violin plots (right) showing the representation of I-A^b^:2W1S tetramer-binding T cells amongst CD4^+^ T cells from the indicated tissues of mice infected with *S.* typhimurium-2W1S versus (*n* = 5). *P* values calculated using one-way ANOVA with Tukey’s multiple comparisons test. **F**: Cumulative violin plots (left) and representative cytometry (right) showing the percentage of CD4^+^ T cells from the indicated tissues that produce IFNγ upon stimulation ex vivo 21 dpi with *S.* typhimurium-2W1S versus control (*n* = 3-6). P values calculated using two-way ANOVA with Bonferroni’s post-test. **G**: Bead-normalised quantification of total dural CD4^+^ T cells expressing IFNy in mice infected orally with *S.* typhimurium-2W1S versus control 21 dpi (*n* = 4-6). P values calculated using two-tailed Student’s unpaired t-test.

### Dural re-location of intestinal CD4 T cells is dependent on CXCR6-CXCL16 interactions

To identify the chemokine-chemokine receptor signals involved in the localisation of intestinal CD4 T cells into the dura, we examined chemokine receptor expression in published scRNA seq data generated from sorted intestinal CD4 T cells^24^. *Cxcr6* was the most highly expressed chemokine receptor in gut CD4 T cells, regardless of the nature of the T cell polarising stimulus ( **Fig. 4A**). We confirmed CXCR6 surface expression on both intestinal and dural Th17 cells following *C. rodentium* infection (**Fig. S5 A-B**). Indeed, the tetramer^+^ CD4 T cells present in the dura following oral challenge with 2W1S-expressing *S.* typhimurium were almost universally CXCR6^+^ (**Fig. 4B**). Within the dura, the CXCR6 ligand, *Cxcl16*, was among the most highly expressed chemokine transcripts (**Fig. 4C**), identifying CXCR6-CXCL16 as a potential axis mediating the recruitment of gut-activated CD4 T cells to the dura. Using our integrated single cell RNAseq dataset of CNS border tissues^25^, we identified meningeal macrophages as the major cell type expressing *Cxcl16*, with more limited expression in cDC and stromal cells (**Fig. 4D**), confirmed at a protein level (**Fig. 4E, S5 C-D**). In contrast to dura, there was limited expression of CXCL16 in splenic macrophages (**Fig. 4F**). To test the importance of this axis in dural CD4 T cell recruitment, we administered a CXCL16-blocking antibody from day 5-17 following 2W1S-*S*. typhimurium challenge (**Fig. 4G**). This led to a reduction in the number of antigen-specific CD4 T cells recruited to the dura (**Fig. 4H**), but not the colon or spleen (**Fig. S5 E**), confirming that the increase in dural CD4 T cells observed following intestinal challenge occurs due to cellular immigration rather than local expansion, and that this recruitment occurs in a Cxcr6-Cxcl16 dependent manner.

**Figure 4.**
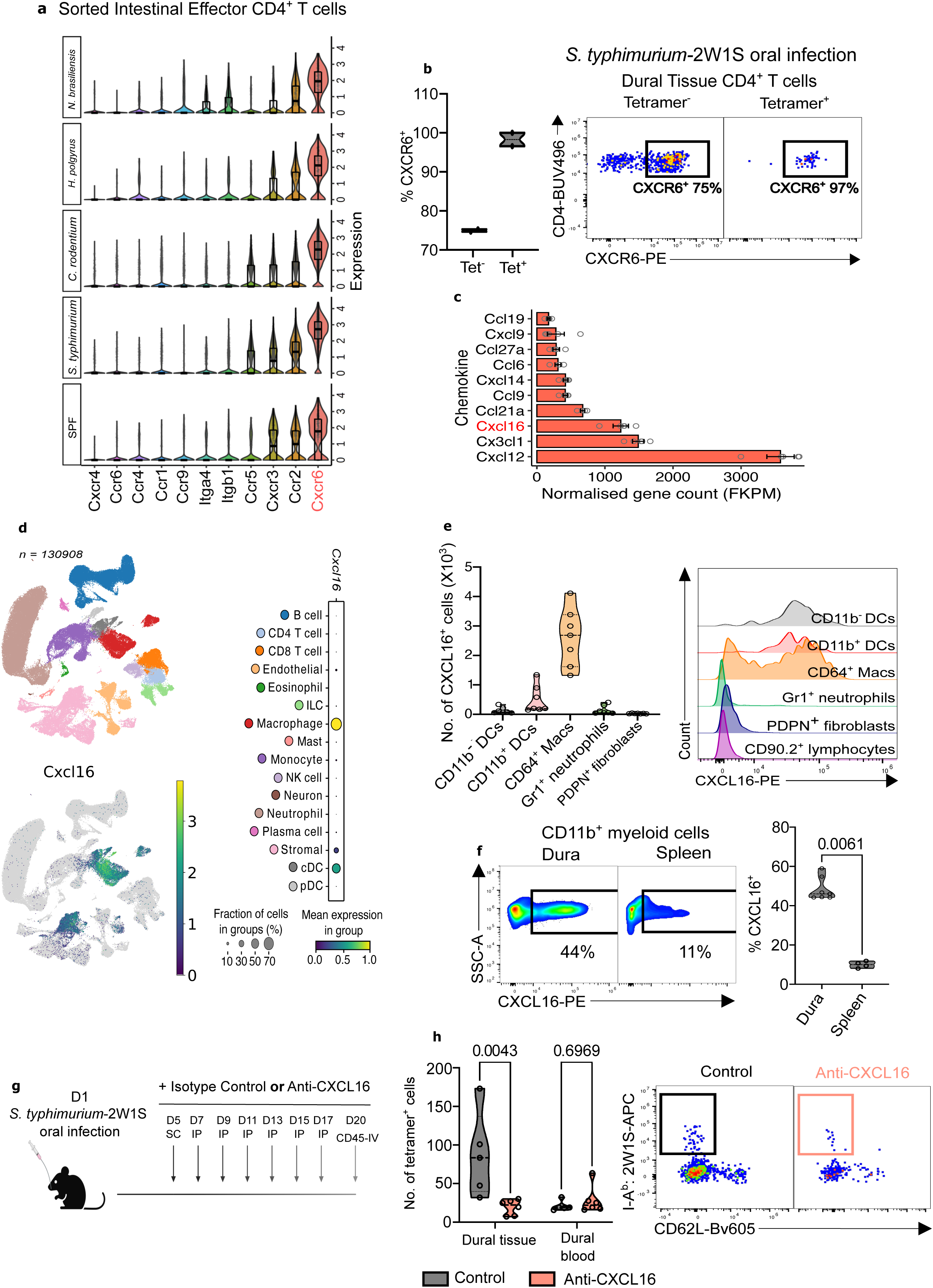
Dural re-location of intestinal CD4 T cells is dependent on CXCR6-CXCL16 interactions **A**: Violin plots showing the top 10 most highly expressed chemokine receptors amongst colonic effector CD4^+^ T cells (*Cd44^+^Sell^-^Ccr7^-^Foxp3^-^*) from mice orally infected with the indicated pathogens. Data derived from published scRNA-seq (Kiner *et. al,* 2021). **B**: Quantification (left) or representative cytometry (right) of CXCR6 expression on dural tetramer^+^ versus tetramer^-^ CD4^+^ T cells in mice infected with 2W1S-expressing *S.* typhimurium (*n* = 2). **C**: Bar plots of the most highly expressed chemokines derived from bulk RNA-seq of whole dural meninges. **D**: (top left) UMAP of 130,908 cells from an integrated scRNA-seq atlas of meningeal tissue in mice (CNSborderatlas.org), colored by cell type. (bottom left) UMAP of *cxcl16* expression or (right) dot plots showing *cxcl16* expression across each cell type. **E**: Cumulative violin plots (left), or cytometry histograms (right) showing cell-specific expression of CXCL16 ex vivo (*n* = 7) **F**: Representative cytometry (left) and cumulative violin plots (right) showing % CXCL16 expression amongst CD11b^+^ cells from indicated organs ex vivo (*n* = 4-7) **G**: Schematic of experimental setup. Mice were orally infected with 2W1S-expressing *S*. typhimurium and received either control isotype antibody or neutralising anti-CXCL16 antibody at the indicated timepoints in the indicated manner (SC = subcutaneous, IP = intraperitoneal, IV = intravenous). **H**: Bead-normalised quantification (left) or representative cytometry (right) showing the relative abundance of dural I-A^b^:2W1S tetramer-binding CD4^+^ T cells following infection with oral *S*. typhimurium-2W1S with anti-CXCL16 treatment versus control (n = 5-6). P values calculated using multiple Mann-Whitney U test with Bonferonni-Dunn multiple comparisons correction.

### Intestinal challenge leads to long-lived dural antigen-specific memory CD4 T cells capable of a recall response upon intravenous re-challenge

Given that gut polarised dural T cells were antigen experienced and displayed an effector memory phenotype (CD62L^low^, CD44^high^), we wondered whether they were capable of expansion following re-challenge with an intravenous pathogen – consistent with our proposal that meningeal immunity is shaped by intestinal cues in order to protect the brain against gut-derived bacteria that enter the blood stream during barrier breach^26^. To test this, we challenged mice orally with 2W1S-*S.* typhimurium, and five and a half weeks later re-challenged intravenously (IV) with the same pathogen (**Fig. 5A**). In mice receiving oral priming followed by an IV re-challenge, there was a significantly higher number of dural 2W1S-specific CD4 T cells at day 5, compared with orally primed mice, whilst tetramer positive cells were almost undetectable in mice receiving an IV challenge alone (**Fig. 5B-C**). Remarkably, these intestinal-derived dural CD4 T cells were long-lived and capable of an antigen-specific recall response following IV re-challenge 5 months after the primary oral challenge (**Fig. 5D**).

**Figure 5.**
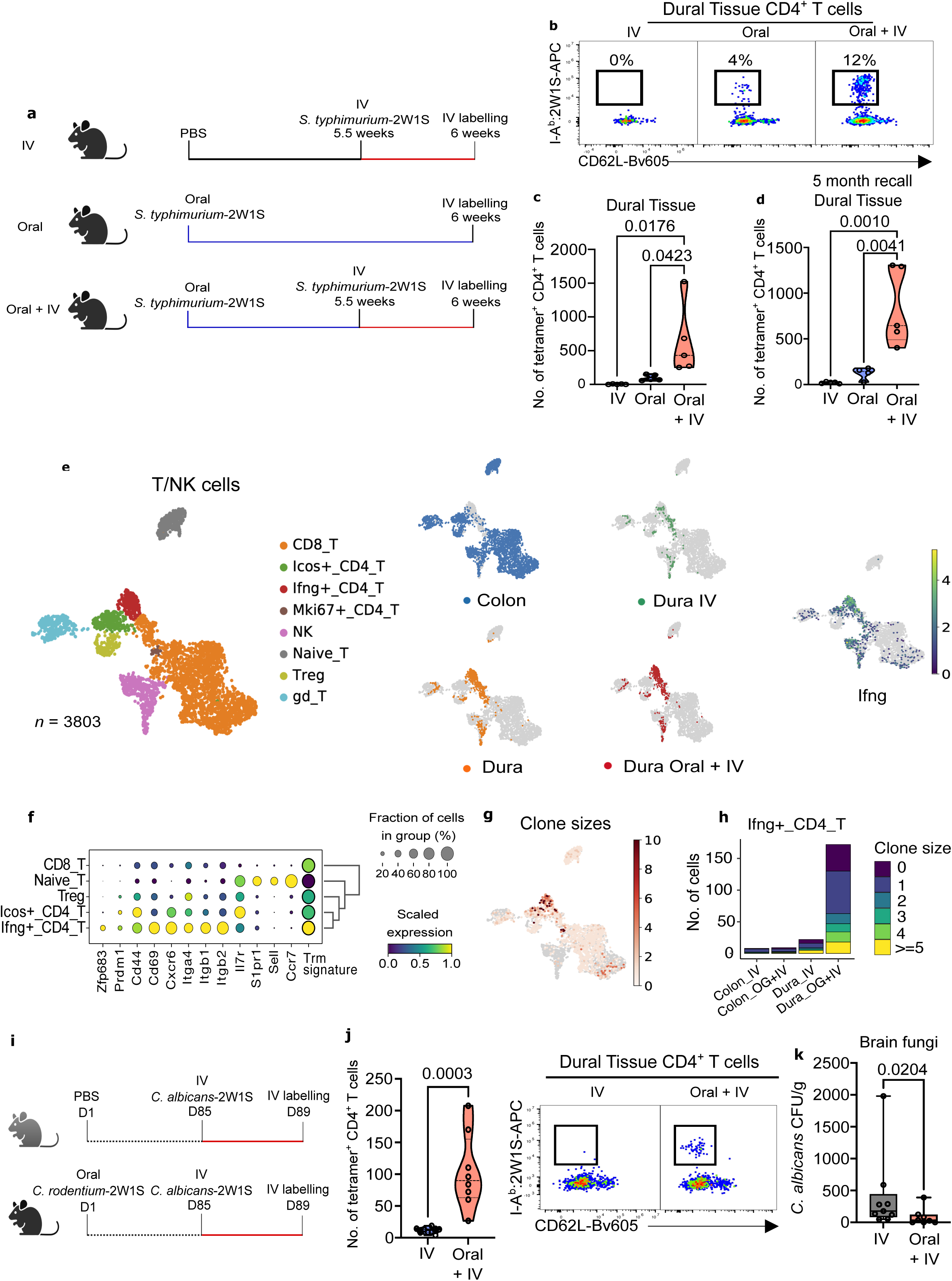
Intestinal challenge generates long-lived dural antigen-specific memory CD4 T cells capable of a recall response upon intravenous re-challenge **A**: Experimental design to detect recall expansion of dural antigen-specific CD4^+^ T cells. Mice were orally gavaged with *S*. typhimurium-2W1S (“Oral” and “Oral + IV”) or PBS control (“IV”). Where indicated at week 5.5, mice received IV injections of *S*. typhimurium-2w1s. 5 days later, mice received IV anti-CD45 and were culled. **B,C**: (b) Representative cytometry or (c) bead-normalised quantification showing the relative abundance of dural I-A^b^:2w1s tetramer-binding CD4^+^ T cells under the indicated conditions (*n* = 5 per group). P values calculated using one-way ANOVA with Tukey’s multiple comparison test. **D:** Bead-normalised quantification of dural I-Ab:2W1S tetramer-binding CD4^+^ T cells when re-challenge with IV S. typhimurium-2W1S was undertaken 5 months rather than 5.5 weeks after oral priming (*n* = 4-5 per group). *P* values calculated using one-way ANOVA with Tukey’s multiple comparison test. **E**: (left) UMAP of 3803 T and NK cells from scRNA-seq of extravascular (IV CD45^-^) CD45^+^ immune cells in the dura or colon, coloured by cell type (left) or (right) by source tissue or treatment arm (for dura only) **F**: Dotplot of selected residency genes and enrichment of tissue-residency lymphocyte signatures amongst CD4^+^ T cells. **G, H:** (g) UMAP of T and NK cells with paired scTCR-seq data coloured by size of expanded T cell clone and (h) bar plot of frequency of cells in the Ifng+_CD4_T cluster by condition, coloured by clone size. **I**: Schematic of experimental layout. Mice were orally gavaged with *C. rodentium-*2W1S (bottom, “Oral + IV”) or PBS control (top, “IV”), and 85 days later intravenously re-challenged with *C. albicans*-2W1S. Mice were culled 4 days later after intravenous labelling with anti-CD45. **J**: Cumulative violin plots (left) and representative cytometry (right) showing the relative abundance of dural I-A^b^:2W1S tetramer-binding CD4^+^ T cells under the indicated conditions (*n* = 8-9). *P* values calculated using Student’s unpaired two-tailed *t-*test **K**: *C. albicans* colony forming units (CFU) per gram of brain tissue between the two groups in (i) (*n* = 8-9). P values calculated using non-parametric Mann-Whitney U test.

Single cell RNA sequencing of extravascular dural and colonic cells following oral priming with *S.* typhimurium followed by IV challenge (oral+IV) or IV challenge alone (**Fig. S6 A-B**) confirmed the presence of an expanded *Ifng*-expressing CD4 T cell cluster, comprised predominantly of cells derived from the dura of oral+IV animals (**Fig. 5E, S7 A-C**). These cells expressed memory and tissue residency markers such as *Cd69*, and showed enrichment of a published lymphocyte tissue-residency gene signature^27^ (**Fig. 5F, S7 D**). They also expressed integrin alpha 4 (*Itga4*) and beta 1 (*Itgb1*), previously shown to be expressed on human memory CD4 T cells in cerebrospinal fluid^28^ and known to be important for lymphocyte entry into the CNS^29^. Expanded TCR clones were largely localised to this *Ifng*^+^ CD4 T cell population, and mainly originated from the oral+IV group (**Fig. 5G-H**), confirming that intestinally primed CD4 T cells localised to the dura are capable of a recall response following IV challenge.

To test whether these dural memory CD4 T cells were functionally important for brain defence, we infected mice orally with a 2W1S-expressing *C. rodentium* to generate dural Th17 cells and 80 days later challenged with an IV 2W1S-expressing *Candida albicans* (**Fig. 5I**). Four days following IV candidal challenge, a robust expansion of dural extravascular 2W1S-specific CD4 T cells was evident in oral *C. rodentium*-primed mice (**Fig. 5J**) and more importantly, reduced fungal load in the brain compared with mice challenged with IV *C. albicans* that had not undergone oral priming (**Fig. 5K, S8 A**). Together, these data confirm the importance of gut activated and polarised, antigen-specific CD4 T cells in CNS defence.

## Discussion

The dural vasculature is fenestrated, enabling bloodborne microbes to access the CNS. In humans, the commonest organisms observed in bacteraemic subjects are gut-derived gram-negative bacteria^30,31^. However, gram negative meningitis is infrequent^32–34^, suggesting that the dura has evolved effective defence mechanisms against gut-derived organisms^26^. We have previously shown that gut educated plasma cells localise to the walls of the dural venous sinuses, providing a defensive shield^4^. Here, we show that there is also an intimate link between gut and dural CD4 T cells, where an infectious challenge in the gut not only activates and polarises CD4 T cells to ensure optimal local antigen-specific responses, but also seeds polarised, antigen-specific memory CD4 T cells to the dura. Here they reside, retaining the capacity to mount a recall response if the same antigen/pathogen is present in the circulation. This link between intestine and dura ensures adaptive cellular immunity in the brain border is tailored to the most likely intravenous pathogen challenge.

Previous studies have described the homeostatic movement of IFNγ-expressing NK cells from the gut to the meninges, promoting the development of anti-inflammatory astrocytes^35^. In addition, CNS perturbations, such as ischaemia, can lead to the induction of IL-17-producing T cells in the small intestine that traffic to the leptomeninges, with deleterious effects on infarct size^36^. Similarly, in experimental autoimmune encephalomyelitis (EAE), pathogenic intestinal-derived CXCR6-expressing Th17 cells were identified in the brain and spinal cord^37^. Here we show that infectious challenges in the gastrointestinal tract can also provide a cue to promote CD4 T cell localisation from the gut to the dura in a CXCR6-CXCL16-dependent manner, and we identify meningeal macrophages as the major source of CXCL16 in the dura.

Meningeal T cells have the capacity to influence brain function via cytokine production. For example, CD4 T cell-derived IFNγ can influence social behaviour^9^, T cell-specific IL-4-deficiency leads to deficits in memory and cognition^10^, and γδ T cell-derived IL-17 promotes synaptic plasticity and short term memory^11^, but in excess can increase anxiety-like behaviour due to direct effects on IL-17 receptor-expressing cortical glutamatergic neurons^12^. A number of studies have also shown that the gut microbiome can influence behaviour and susceptibility to anxiety and depression, including via IL-17^38–41^. The assumption had been that these effects on brain function were due to the impact of increased circulating cytokines produced by Th17 cells in the gut^41^. Our data provide an alternative explanation, showing that cytokine-producing CD4 T cells activated by gut microbes may localise to the dura. We also found that during DSS colitis, a model of inflammatory bowel disease (IBD), there was an expansion of Th17 cells in the dura. This observation has important implications for the mechanistic underpinning of the increase in mental health problems described in patients with IBD, with up to 30% and 25% of patients experiencing anxiety and depression symptoms respectively^42^. Our work raises the possibility that gut-derived IL-17-secreting CD4 T cells may contribute to this phenomenon.

Schistosomiasis affects 140 million people world-wide and is prevalent in poor communities without clean water in tropical and sub-tropical areas, particularly in sub-Saharan Africa^43^. Here we show that *S. mansoni* infection is associated with a substantial expansion in dural Th2 cells, with important clinical implications. Although IL-4 deficiency has been associated with deficits in memory and learning, the effect of a chronic excess of IL-4 in the brain has not been assessed and could contribute to the epidemiological associations observed between chronic schistosomiasis and academic attainment that were first described more than 70 years ago^44,45^.

More broadly, our work has implications for our understanding of how the brain might sense systemic perturbations. We have long known that sensory neurons provide a direct means by which information from peripheral organs, including the gut, can be conveyed to the brain. Our findings here that CD4 T cells, and previously B/plasma cells^4^, can migrate to the dura provide an additional mechanism by which information can be transmitted to the CNS, utilising the physical re-location of immune cells that communicate with neurons via cytokines, with cytokine-receptor expression increasingly recognised on cells of the CNS^9,46–49^. This provides an information relay system distinct from neuronal signalling, employing migratory lymphocytes rather than axons that physically span the distance between peripheral organs and the brain, and cytokines rather than neurotransmitters to transfer signals from cell to cell. Ultimately, this link between intestinal and dural immune cells allows the CNS to accrue immunological memory of microbes most likely to enter the blood and compromise organismal survival.

In summary, our data shows that gut challenge establishes and shapes long-lived CD4 memory in the dura to protect CNS organs from bloodborne pathogens, with the potential to influence brain function and behaviour via local cytokine production.

## Methods

### Mice

Mouse experiments were carried out in specific pathogen-free conditions at the University of Cambridge and Pathogen Genomics, Wellcome Sanger Institute under relevant project licences with institutional study plan approval by a local Animal Welfare and Ethical Review Board (AWERB) in accordance with the United Kingdom Animals (Scientific Procedures) Act of 1986. Husbandry and animal welfare was provided daily by animal technicians. Mice were given *ad libitum* access to food (Safe Diets, Safe) and water. Following intravenous (IV) infections, mice were additionally given Pure-Water Gel:Total Hydration (Bio-Serv, PWG-2) and a powdered version of the above food mixed with water in petri dishes on their cage floor daily.

C57BL/6J wild-type (WT) mice were used unless otherwise specified. For. *S. mansoni* infections, TO outbred (HsdOla:TO) or BALB/C mice (bred in house at the Sanger Institute) were used for flow cytometry or imaging, respectively under the licence of Gabriel Rinaldi (P77E8A062). WT mice were bred in-house or purchased from Jackson Laboratories (Margate, UK) and acclimatised for at least one week pre-experimentation. RORγt-eGFP reporter mice were kindly provided by Andrew McKenzie (MRC-LMB, UK).

Where specified, mice received IV anti-CD45 antibody injections (1.5 μg per mouse in 100 μL sterile PBS) 3 minutes before sacrifice. Mice were culled by CO_2_ exposure followed by either cervical dislocation or exsanguination. Peripheral blood was taken to confirm successful blood CD45 labelling.

### *Ex-vivo* stimulation of cells for cytokine/chemokine detection

For cytokine/chemokine detection, colonic and splenic immune cells were stimulated as cellular suspensions, whereas dura was stimulated as whole tissue prior to digestion/processing. Cytokine stimulation was undertaken at 37 °C/5% CO_2_ for 4 hrs in complete RPMI (RPMI plus 10% FCS and 10 mM HEPES) plus phorbol 12-myristate 13-acetete (PMA) (20 ng/mL, Sigma-Aldrich, P1389), ionomycin (1 μg/mL, Sigma-Aldrich, I9657) and BD GolgiPlug protein transport inhibitor (1:1000, 555029, BD Biosciences). Chemokine stimulation was undertaken under the same conditions, but without PMA or ionomycin.

### *S*. typhimurium oral and intravenous infections

Frozen glycerol (20% v/v) stocks of *S. enterica* servovar typhimurium-2W1S (*S.* typhimurium) strain BRD509^50^ were defrosted at room temperature (RT), streaked onto agar plates and incubated upside down overnight at 37 °C. 3-4 colonies were added to 100 mL LB broth and incubated at 37 °C overnight with no shaking. Cultures were diluted in PBS and mice either received 200 μL (1 × 10^7^ CFUs) of inoculum by oral gavage, or 100 μL (1 × 10^6^ CFUs) by tail-vein injection. Infection CFU counts were verified by streaking the inoculums onto agar plates containing 100 μg/mL streptomycin and incubating overnight at 37 °C. 24 hours prior to oral infection, mice were gavaged with 20 mg of streptomycin (Bio Basic, SB0494) in 100 μL PBS to aid colonisation. Faecal CFUs were plated 4-10 days post-oral gavage to confirm stable colonisation of mice. Mice were intravascularly labelled with IV anti-CD45 prior to culling.

### *C. rodentium* oral infections

Frozen 20% glycerol (v/v) stocks of *C. rodentium* (strain ICC180) were defrosted at RT, streaked onto agar plates and incubated upside down overnight at 37 °C. 3-4 colonies were added to 100 mL LB broth and incubated at 37 °C overnight with shaking (180 rpm). The culture was spun down and concentrated in sterile PBS to yield 1 × 10^9^ CFUs per 200 μL inoculum for oral gavage. Inoculums were plated immediately post-infection on agar plates at 37 °C overnight to confirm infection CFU counts. Faecal CFUs were plated 4-10 days post-oral gavage to confirm stable colonisation of mice.

### Construction of *C. rodentium* expressing 2W1S

The 2W1S peptide was inserted between positions 126 and 127 of the *C. rodentium* EspA translocator protein in strain ICC169. The homology regions flanking the insertion site of espA, comprising the upstream and downstream 300 bp, and the intertwine sequence of the peptide, were synthesized (Thermo) and cloned into the SacI-SphI sites of pSEVA612S^51^. The resulting pSEVA612S-2WtagEspA was transformed into E. coli CC118λpir^52^. The endogenous *C. rodentium espA* gene was then replaced with 2WtagEspA via tri-parental conjugation as described^53^. All the plasmids and mutants were confirmed by sequencing (Eurofins).

### *S. mansoni* life cycle maintenance and infections

The life cycle of the NMRI (Puerto Rican) strain of *S. mansoni* was maintained by infection of TO outbred (HsdOla:TO, Inotiv, UK) female mice and Biomphlaria glabrata snails (bred and maintained at the Sanger Institute). Cercariae were collected from infected snails exposed to bright light and counted under a dissecting microscope. Mice were anaesthetised with isofluorane and infected percutaneously via their tail, which was sub-merged into a tube containing 500 μL of a stock solution containing 600 cercariae/mL.

Mice were culled 42 days post-infection by intraperitoneal injection of 200 mg/mL pentobarbital supplemented with 100 U/mL heparin. Mice were intracardially perfused with phenol red-free DMEM supplemented with 10 U/mL heparin and liver tissue collected and digested to isolate eggs to maintain the lifecycle infection of the snail.

### *T. muris* infections

Mice were infected by oral gavage with a low (20-25) dose of embryonated eggs from *T. muris* (eggs were obtained from Prof. R. Grencis) and culled at Day 21 after intravascular labelling with IV anti-CD45.

### *C. albicans* intravenous infections

Frozen glycerol (20%, v/v) stocks of *C. albicans*-2w1s were defrosted at RT, streaked onto yeast extract peptone dextrose (YPD) plates (2% peptone, 2% glucose, 1% yeast extract, 2% agar, 1% penicillin/streptomycin), and incubated upside down overnight at 30 °C, 5% CO_2_. 3-4 colonies were added to 50 mL YPD broth (2% peptone, 2% glucose, 1% yeast extract) and shaken at 180 rpm for 18 hours at 30 °C, 5% CO_2_. Yeast cells were washed in PBS and enumerated under light microscopy to generate an inoculum of 1 × 10^5^ CFUs in 100 μL PBS. Inoculums were plated immediately post-infection on YPD plates at 30 °C overnight to confirm exact infection CFU counts. Mice were intravascularly labelled with IV anti-CD45 prior to culling.

To enumerate *C. albicans* CFUs in brain, a single cerebral hemisphere was weighed and homogenised with 500μL PBS in a 2 mL Eppendorf tube using a 1 mL plastic syringe. Homogenates were centrifuged to remove large debris and 100 μL of homogenate was added to a YPD plate and spread using glass beads. Plates were incubated upside down at 30 °C, 5% CO_2_ for 48 hours and visible colonies counted and normalised to tissue mass.

### CXCL16 neutralisation

5 days following oral gavage with *S.* typhimurium-2W1S, mice sedated with isoflurane received a subcutaneous scalp injection of 200 μL anti-CXCL16 (1 mg/mL, BioTechne, MAB503) or Rat IgG2a control (1 mg/mL, BioTechne, MAB006) reconstituted in carboxymethylcellulose hydrogel (10 mg/mL in sterile PBS, Sigma-Aldrich, C9481). Mice subsequently received 100 μL of the specified antibodies diluted to the same concentration in sterile PBS by intraperitoneal injection every second day until Day 17. Mice were culled on Day 20 after intravascular labelling with anti-CD45. Treatment groups were equally split between cages to minimize cage effects.

### DSS-induced colitis

For flow cytometry experiments, drinking water was supplemented with 2.5% (w/v) dextran sulfate sodium (DSS, MP Biomedicals) for 5 days, followed by 14 days on normal drinking water, a further 5 days on 2.5% DSS and a final 6 days on normal drinking. Mice were culled after intravascular labelling with anti-CD45.

For TCR sequencing experiments, mice were given 2.5% (w/v) DSS in drinking water for 5 days, followed by 16 days on normal drinking water prior to culling and perfusion.

### Colon tissue processing

Extraneous fat was removed using curved scissors and faecal content extruded using forceps. The colon was opened longitudinally, washed in ice-cold PBS/HEPES, and cut into 0.5 cm pieces for transfer into epithelial stripping buffer (5 mL RPMI, 2% FCS, 10 mM HEPES, 5 mM EDTA, 1 mM DTT). This was incubated at 37 °C shaking for 25 min, vortexed, and washed in PBS/HEPES through a wire-mesh filter before transfer into digest buffer (5 mL RPMI, 2% FCS, 10 mM HEPES, 1 mg/mL collagenase I [Thermo Fisher Scientific, Cat. No 17100017], 100 μg/mL DNAse I [Merck, Cat. No 10104159001]). Digestion was performed with shaking at 37 °C for 30-40 min followed by vortex. Homogenates were passed through a 70 μm cell filter, mashed using the rubber end of a 1 mL syringe insert, and washed with 3% FCS/PBS. Immune cell-enriched suspensions were obtained via percoll-density centrifugation (20 mins, 600 x g). Cells were harvested from the 40:80% interface.

### Dura tissue processing

Skull cap dura was dissected in ice-cold PBS under a dissecting microscope, transferred to 1 mL digest medium (1 mL RPMI, 2% FCS, 10 mM HEPES, 2.5 mg/mL Collagenase D [Thermo Fisher Scientific, 11088866001], 125 μg/mL DNAse I [Merck, 10104159001]), and incubated for 30 min at 37 °C with gentle agitation at 15 min. Tissue was vortexed vigorously, pipetted up and down, and passed through a 70 μm FACS tube filter.

### Spleen tissue processing

Spleens were mechanically disaggregated through a 70 μm cell filter using the rubber end of a 1 mL syringe insert. Cells were washed with 3% PBS/FCS and resuspended in 2 mL RBC lysis buffer for 1 min. Lysis was quenched with 3% PBS/FCS. 1/10 of spleen samples were used for flow cytometry staining.

### Flow cytometry

Staining was carried out in 96 well V-bottom plates (Thermo Scientific Fisher, 612V96). Precision Count Beads (BioLegend, 424902) were added to enumerate total cell number. Live:dead discrimination was achieved using ViaKrome 808 Fixable Viability Dye (Beckmann Coulter, C36628) or Live/Dead Near-IR (for cell sorting), and blocking was performed using 3% PBS/FCS containing 0.5% (v/v) heat-inactivated normal mouse/rat serum (Invitrogen, 10410/Thermo Scientific Fisher, 10-710-C).

2w1s (EAWGALANWAVDSA) peptide-loaded mouse I-A(b) tetramers conjugated to APC or PE were provided by the NIH Tetramer Facility, Atlanta, Georgia. These were incubated with cells in PBS at RT for 1 hour.

Extracellular staining was carried out in 3% PBS/FCS on ice for 30 mins. Cells were fixed for 4 mins at RT in 4% paraformaldehyde and washed in 3% PBS/FCS before analysis.

Intracellular staining was carried out according to manufacturer’s instructions with CytoFix/CytoPerm (BD Biosciences, 554714) used for cytokine/chemokine detection, and the Foxp3 Transcription Factor Staining Kit (eBioscience, 11500597) used for transcription factors.

Flow cytometry was performed on a CytoFLEX LX Flow Cytometer (Beckman Coulter) at the Core Flow Facility at the Laboratory of Molecular Biology. Cell sorting was performed on a BD Influx. Antibodies for cell staining are as follows (clone/cat. No/dilution): anti-IFN-γ-FITC (XMG1.2, 505806 BioLegend, 1:200), anti-CD45 FITC (30-F11, 103108 BioLegend, 1:200), anti-CD11c FITC (N418, 35-0114 eBioscience, 1:200), anti-CD11b FITC (M1/70, 101206 BioLegend, 1:300), anti-CD45 BUV395 (30 F-11, 564279 BD Biosciences, 1:200), anti-CD4 BUV496 (GK1.5, 612952 BD Biosciences, 1:200), anti-CD19 PerCp (6D5, 115532 BioLegend, 1:200), anti-B220 PerCp-Cy5.5 (RA3-6B2, 103236 BD Biosciences, 1:200), anti-MHC-II eFlour450 (AF6-120.1, 48-5320-82 eBioscience, 1:400), anti-CD45 PacBlue (30-F11, 103126 BioLegend, 1:200), anti-TCRβ Bv421 (H57-597, 109230 BioLegend, 1:100), anti-CD90.2 Bv510 (53-2.1, 140319 BioLegend 1:400), anti-CD3ε Bv605 (145-2C11, 100351 BioLegend, 1:100), anti-CD62L Bv605 (MEL-14, 104437 BioLegend, 1:200), anti-CD62L Bv786 (MEL-14, 564109 BioLegend, 1:200), anti-Gr1 Bv786 (RB6-8C5, 740850 BD Biosciences, 1:400), anti-GATA3 PE (TWAJ, 12-9966-42, eBiosciences, 1:200), anti-IL-17a PE (eBio17B7, 12-7177-81 eBiosciences, 1:200), anti-T-bet PE (eBio4B10, 12-5825-82, eBiosciences, 1:200), anti-CXCL16 PE (12-81, 566740 BD Biosciences, 1:200), anti-CXCR6 PE (SA051D1, 151103 BioLegend, 1:200), anti-TCRγδ PE-Cy7 (GL3, 118124 BioLegend, 1:200), anti-Podoplanin PE-Cy7 (8.1.1, 127411 BioLegend, 1:200), anti-RORγt APC (B2D, 17-6981-82 eBiosciences, 1:200), anti-GATA-3 (TWAJ, 50-9966-42 eBioscience, 1:200), anti-CD64 APC (X54-5/7.1, 139306 BioLegend, 1:200), anti-CD45 APC (QA17A26, 157605 BioLegend, 1:200) anti-CD8 APC-Cy7 (YTS156.7.7, 126620 BioLegend, 1:200).

### Confocal imaging and staining

Mice were intracardially perfused with 20 mL ice-cold PBS and skull caps fixed in Antigenfix (P0016, Diapath) for 90 mins on ice, washed in PBS, and kept in PBS/FCS until staining was performed. Dura was blocked at RT for 30 minutes in blocking-permeabilization solution (2% normal rat serum, 2% normal mouse serum, 0.1% Triton-X-100, 0.05% Tween and 0.1% saponin in 0.1 M TRIS) and stained for antigens overnight at 4 °C using antibodies in the same blocking solution. Primary stains included rabbit anti-GATA-3 (EPR16651, ab199428 Abcam, 1:50), anti-CD31 AlexaFlour647 (MEC13.3, 102516 BioLegend, 1:100), anti-CD4 PE (GK1.15, 12-0041-82 ThermoFisher Scientific, 1:100), and rabbit anti-GFP AlexaFlour 488 (polyclonal, A2131 ThermoFisher Scientific, 1:100). Secondary stains used donkey anti-rabbit IgG AlexaFlour 488 (A32790, ThermoFisher Scientific, 1:250). Three 30 minute washes at RT were performed with PBS between and after the two rounds of staining. Dura was mounted in Flouromount-G (0100-01, SouthernBiotech) and coverslips applied for imaging. Imaging was carried out using a Zeiss LSM 880 confocal microscope in super-resolution Airyscan mode with image analysis carried out in Zeiss ZEN blue.

### Bulk TCR sequencing sample preparation

Perfused dura and intestinal tissues were placed in RNA Later (AM7020, ThermoFisher Scientific) until RNA extraction with the RNeasy plus micro kit (Qiagen) according to manufacturer’s instructions. Libraries were prepared using SMARTer Mouse TCR a/b Profiling Kit (Takara). Libraries were pooled at equimolar concentration and sequenced on MiSeq (Illumina) on 2 × 300bp run. Resulting libraries were demultiplexed using Casava (Illumina).

### Bulk TCR sequencing analysis

Fastq files were processed using mixcr (v3.0.13) with the ‘analyze amplicon’ workflow with default settings (--species mmu --starting-material rna --5-end v-primers --3-end j-primers--adapters adapters-present --receptor-type tcr). The output files were then converted using vdjtools (v1.2.1). For each sample, clonotypes were grouped by according to whether sequences displayed identical V and J gene usage and CDR3 amino acid sequences. The clonotypes were then tabulated per treatment per animal per tissue source per locus (TRB or TRA). To tabulate clonotype overlap/sharing, samples were randomly down sampled to the same sequencing depth (100000 total counts) and the mean percentage overlap (relative to total meningeal clonotype counts) was tabulated over 100 iterations.

### Bulk RNA-seq library construction and sequencing

Perfused dura was placed in RNA Later (AM7020, ThermoFisher Scientific) until RNA extraction with the RNeasy plus micro kit (Qiagen) according to manufacturer’s instructions. Genomic DNA contamination was removed using Optimal DNA depletion columns (Qiagen). The quality and concentration of purified RNA was assessed on Bioanalyzer 2000 (Applied Biosystems). Libraries were prepared using the SMARTer stranded total RNAseq mammalian pico input kit (Takara) according to the manufacturer’s instructions. 5 ng of total RNA was used for production of libraries. Resulting libraries were reassessed on Bioanalyzer 2000. Libraries were pooled at an equimolar concentration. Sequencing was performed using Hiseq 4000 (Illumina) on a 2 x 150 bp sequencing run. Pooled libraries were demultiplexed using Casava (Illumina).

### Bulk RNA-sequencing processing and analysis

Fastq files from sequencing libraries were trimmed of the first 3 nucleotides of the R1 strand. Contaminating adaptor sequences and poor-quality bases were removed using Trim Galore (Babraham bioinformatics). Libraries were only trimmed for quality. Quality of resulting files were assessed by FastQC and aligned to the mm10 genome using HISAT2, followed by gene level quantification using the Featurecount function from the Rsubread package in R. All subsequent analysis was performed in R. For within-sample comparisons of chemokine expression, FPKM normalisation was applied.

### Single cell library construction and sequencing

Gene expression libraries were prepared from FACS-sorted populations of single cells as specified in the figures using the 10x Chromium Controller and 10x Chromium Single Cell V(D)J 5’ kit v2 with TCR amplification (10x Genomics, Inc.). Samples were loaded onto the chip following manufacturer’s recommendations. The resulting library was sequenced on the Illumina NovaSeq 6000 platform (paired-end sequencing, 2 x 150 bp). BCL files were demultiplexed using Casava (Illumina). The resulting fastq files were assessed using FastQC (v0.11), FastqScreen (v0.9) and FastqStrand (v0.0.5) prior to alignment (mm10) and downstream processing with the CellRanger (v7.0) pipeline.

### Processing of scRNA-seq

Single-cell gene expression data from CellRanger count outputs (filtered features, barcodes, and count matrices) were analysed using the Scanpy^54^ (v1.9.1) workflow. Doublet detection was performed using Scrublet^55^ (v0.2.1), with cells from iterative sub-clustering flagged with outlier Scrublet scores labelled as potential doublets. Cells with counts mapped to >8000 or <200 genes were filtered. The percentage mitochondrial content cut-off was set at <10.0%. Genes detected in fewer than 3 cells were filtered. Total gene counts for each cell were normalised to a target sum of 104 and log1p transformed. This resulted in a working dataset of 11,655 cells. Highly variable features were selected based on a minimum and maximum mean expression of ≥0.0125 and ≤3 respectively, with a minimum dispersion of 0.5. Total feature counts, mitochondrial percentage, and cell cycle (G2M, S phase) scores were regressed before scaling. The number of principal components used for neighbourhood graph construction was set to 30 initially, and subsequently 20 for subgroup analysis. Clustering was performed using the Leiden algorithm with resolution set at 1.0. Uniform manifold approximation and projection (UMAP, v0.5.1) was used for dimensional reduction and visualisation, with a minimum distance of 0.3, and all other parameters according to the default settings in Scanpy.

### Analysis of scRNA-seq

Cell types of interest were subset and re-clustered iteratively as described above. Resulting clusters were annotated using canonical marker genes and top differentially expressed genes. Differential gene testing was performed using the Wilcoxon rank sum test implemented in Scanpy’s tl.rank_genes_groups. Gene set scoring was performed using Scanpy’s tl.score_genes tool. The gene set representing a tissue resident lymphocyte signature was obtained from a published RNAseq dataset^27^. Gene lists for chemokines or chemokine receptors were obtained from the gene ontology (molecular function) database. Gaussian kernel density estimation was performed using Scanpy’s tl.embedding_density function. Unless otherwise indicated, log-transformed expression values were used for plotting. For assessment of *Cxcl16* expression in dural tissues, CNSbordercellatlas.org, an integrated atlas of published scRNA-seq data of the meninges, was downloaded and analysed^25^. For assessment of chemokine receptor expression on CD4^+^ T cells in the gut, scRNA-seq of sorted colonic CD4^+^ T cells was obtained from GSE160055 and processed and annotated according to the authors’ descriptions^24^. All publicly available data was processed and analysed in Scanpy, as described above (v1.9.1).

### Processing and analysis of scTCR-seq

Sequenced single-cell V(D)J (TCR) libraries were first processed with CellRanger vdj and resulting all_contig.fasta and all_contig_annotations.csv files were re-annotated and processed using the dandelion (v0.2.5) pipeline^56^. TCR contigs were filtered according to cell droplets that passed QC in the scRNA-seq GEX data and non-productive TCR contigs were removed. TCR sequences were assigned to single-cell GEX profiles based on their droplet barcode. Clonal V(D)J clustering was performed based on heavy-and light-chain V-J sequences and junctional sequences. Resulting data was visualised in Scanpy (v1.9.1).

## Data availability

Sequencing data have been made available on the GEO public repository under accension numbers GSE233489, GSE233490, GSE233491. Published scRNA-seq data and accompanying metadata were accessed and downloaded from the GEO repository using the ascension GSE160055.

## Acknowledgements

A.F. was supported by a Wellcome Trust PhD Studentship (102163/B/13/Z); DP is supported by a National Council of Science and Technology (CONACYT) of Mexico and Cambridge Trust PhD scholarship; C.Y.L by a Gates Scholarship; A.S. by an NIHR Clinical Lecturership, Z.K.T. and M.R.C. by a Medical Research Council Research Project grant (MR/S035842/1); D.R.W. by a Senior Research Fellowship from the Wellcome Trust 110199/Z/15/Z; M.R.C. and R.D.M.B were supported by an NIHR Research Professorship (RP-2017-08-ST2-002), and M.R.C, A.P. and M.C. by a Wellcome Trust Investigator Award (220268/Z/20/Z). G.F. and D.R.-G. were supported by a Wellcome Trust Investigator Award (107057/Z/15/Z). S.C. and KH were supported by the Wellcome Trust (098051). The Clatworthy lab utilises infrastructure supported by the National Institute of Health Research (NIHR) Cambridge Biomedical Research Centre (NIHR203312). We thank Professor S. McSorley (Salmonella-2W1S) for provision of bacterial strain. We are grateful for the tetramers produced and provided by the NIH Tetramer Facility, Atlanta, Georgia and for the use of the Core Flow Cytometry Facility at the Laboratory of Molecular Biology. The views expressed are those of the authors and not necessarily those of the NIHR or the Department of Health and Social Care.

---

**Supplemental Figure 1.**
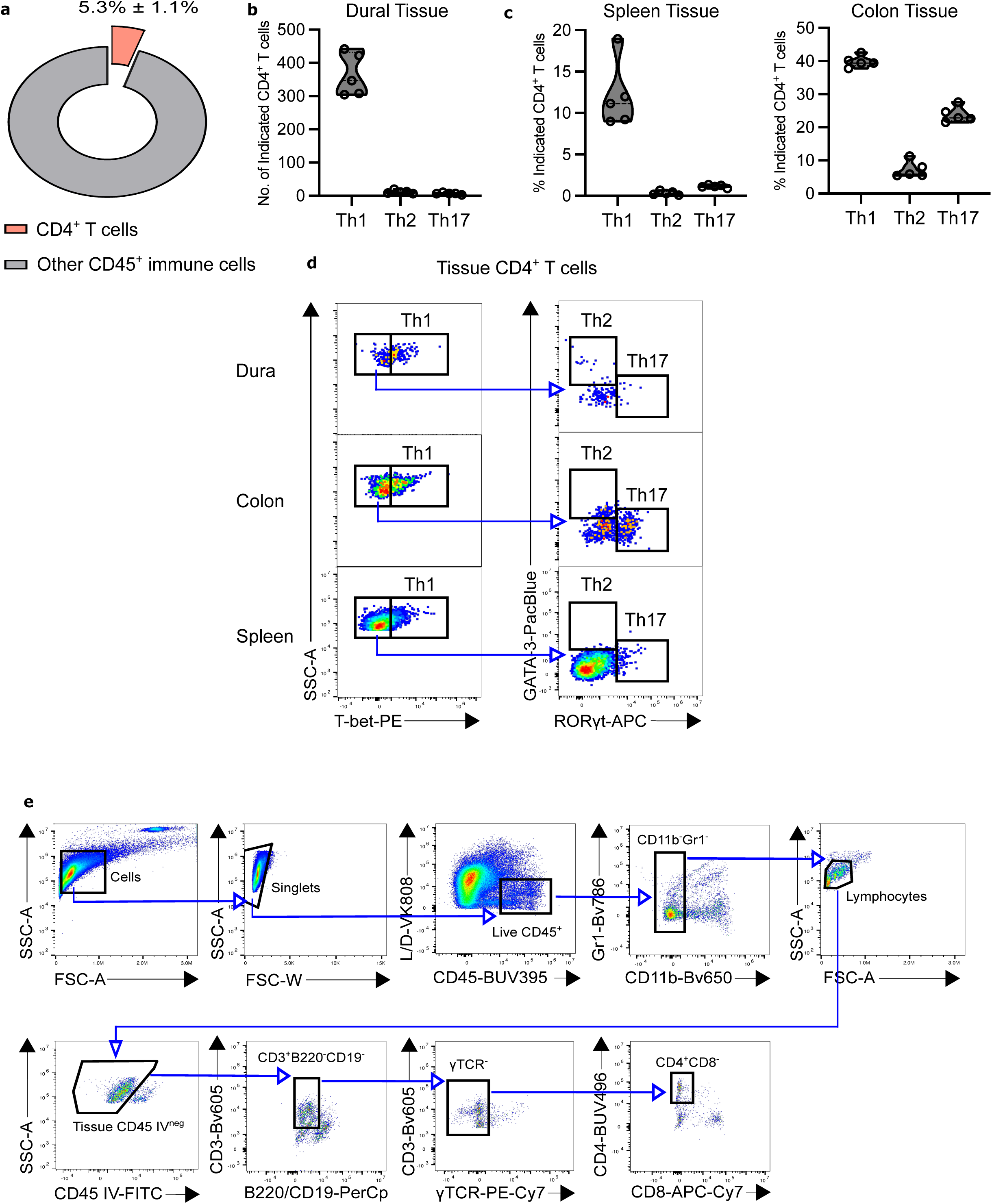
(related to Main Figure 1 A) **A**: Representation of dural CD4^+^ T cells (CD45IV^neg^) amongst total CD45^+^ dural immune cells **B**: Bead-normalised quantification of the indicated dural Th cell subsets (*n* = 5) **C**: Representation of Th cell subsets amongst CD4^+^ T cells from spleen (left) or colon (right) (*n* = 5) **D**: Representative cytometry of CD4^+^ Th cell subsets from the indicated organs. **E**: Representative gating strategy to identify dural CD4^+^ T cell subsets.

**Supplemental Figure 2.**
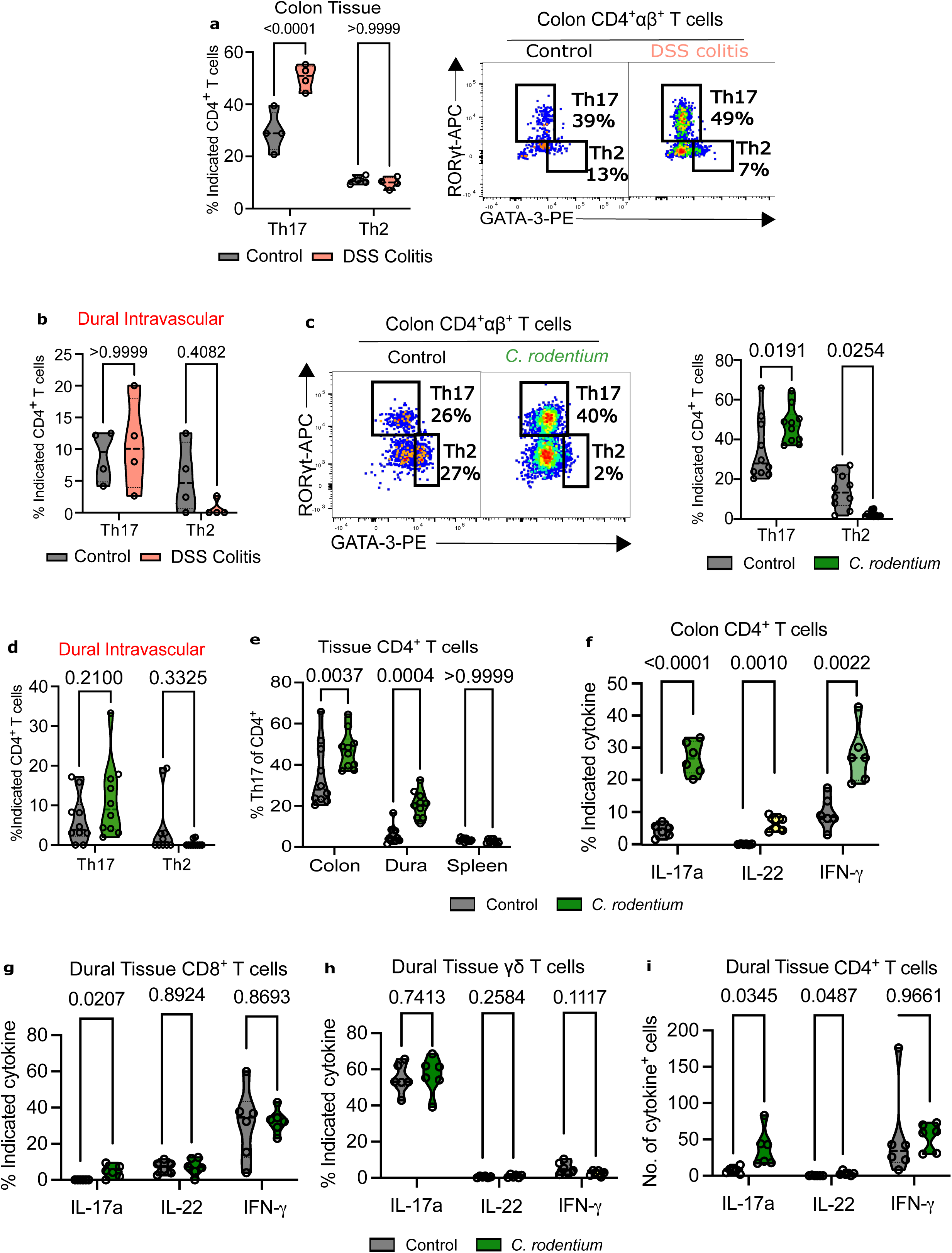
(related to Main Figure 1 B-J) **A:** Cumulative violin plots (left) and representative cytometry (right) of colonic Th17 and Th2 cells in DSS colitis *versus* control (*n* = 4). **B**: Cumulative violin plots of dural intravascular (IV-CD45^+^) Th17 and Th2 cells in DSS colitis *versus* control (*n* = 4). **C**: Cumulative violin plots (right) and representative cytometry (left) of colonic Th17 and Th2 cells in *C. rodentium*-infected mice (21 dpi) *versus control* (*n* = 10). **D**: Cumulative violin plots of dural intravascular (IV-CD45+) Th17 and Th2 cells in *C. rodentium* infection *versus* control (*n* = 10). **E**: Representation of Th17 cells amongst total CD4^+^ T cells from the indicated organs in *C. rodentium*-infected mice (21 dpi) *versus* control (*n* = 4). **F**: Percentage of colonic CD4^+^ T cells from *C. rodentium*-infected *versus* control mice expressing the indicated cytokine upon stimulation *ex vivo* (*n* = 6). *P* values calculated using multiple unpaired student *t*-test Bonferroni-Dunn multiple comparisons correction. **G**: Percentage of dural CD4^+^ T cells from *C. rodentium*-infected *versus* control mice expressing the indicated cytokine upon stimulation *ex vivo* (*n* = 6) mice. *P* values calculated using multiple unpaired student t-test and Bonferroni-Dunn multiple comparisons correction **H**: Percentage of dural γδ T cells from *C. rodentium*-infected *versus* control mice expressing the indicated cytokine upon stimulation *ex vivo* (*n* = 6). *P* values calculated using multiple unpaired student *t*-test and Bonferroni-Dunn multiple comparisons correction **I**: Bead-normalised quantification of total cytokine-expressing dural CD4^+^ T cells from *C. rodentium*-infected *versus* control mice (*n* = 6). *P* values calculated using multiple unpaired student t-test and Bonferroni-Dunn multiple comparisons correction. *P* values derived by two-way ANOVA with Bonferonni’s post-test (A, B, C, D) or multiple unpaired student’s *t*-test with Bonferroni-Dunn multiple comparisons correction

**Supplemental Figure 3.**
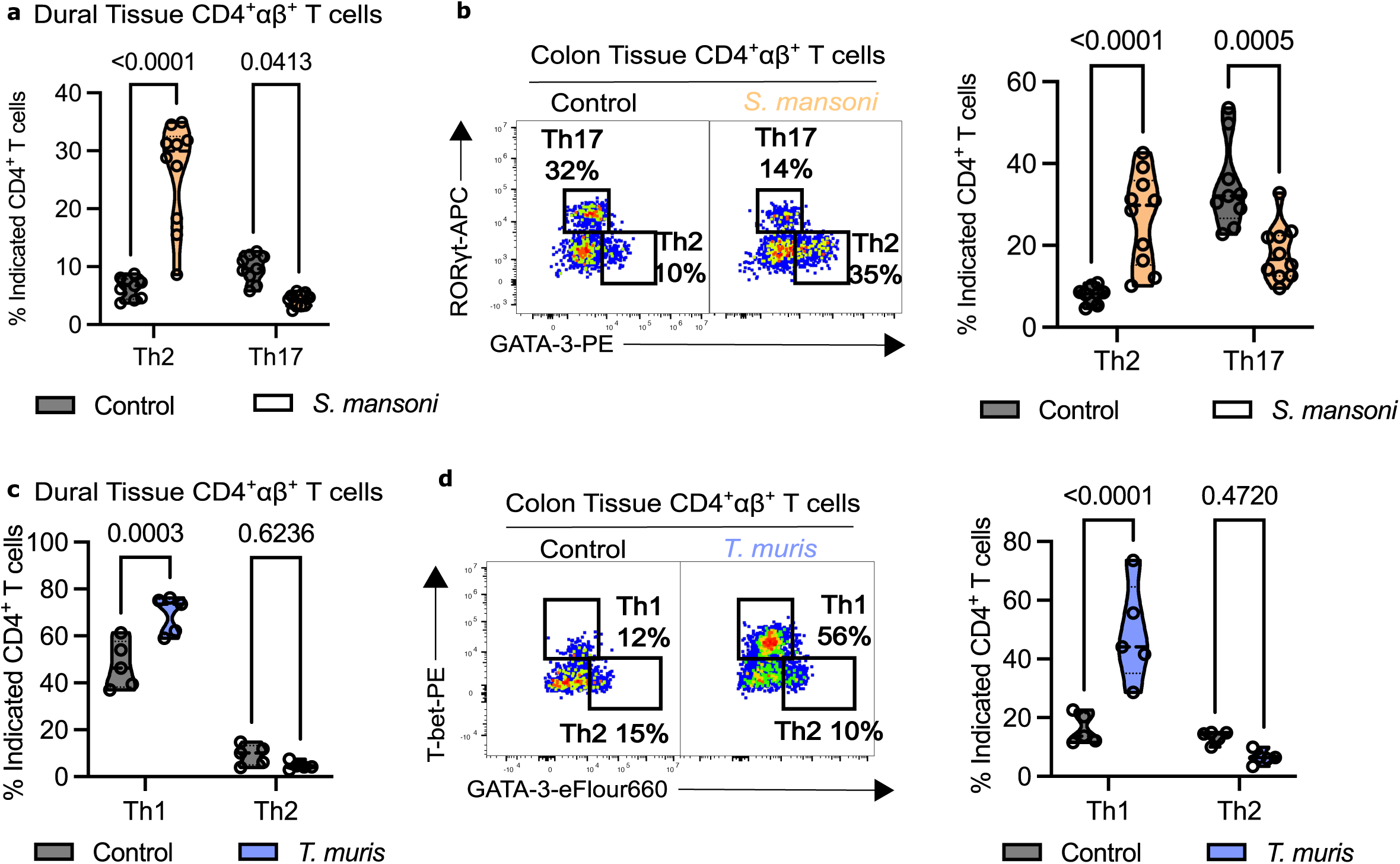
(related to Main Figure 2 A-B, D-E) **A:** Representation of dural Th2 and Th17 cells from *S.mansoni*-infected versus control mice (42 dpi). (*n* = 9-10). **B**: Representative cytometry (left) and cumulative violin plots (right) showing the representation of Th17 and Th2 cells amongst total colonic CD4^+^ T cells from *S.mansoni*-infected *versus* control mice (42 dpi). (*n* = 9-10). **C**: Representation of Th1 and Th2 cells amongst total dural CD4+ T cells from *T. muris*-infected versus control mice (21 dpi). (*n* = 5). **D**: Representative cytometry (left) and cumulative violin plots (right) showing the representation of Th17 and Th2 cells amongst total colonic CD4^+^ T cells from *T. muris*-infected versus control mice (21 dpi). (*n* = 5). *P* values derived using two-way ANOVA with Bonferroni’s post-test.

**Supplemental Figure 4.**
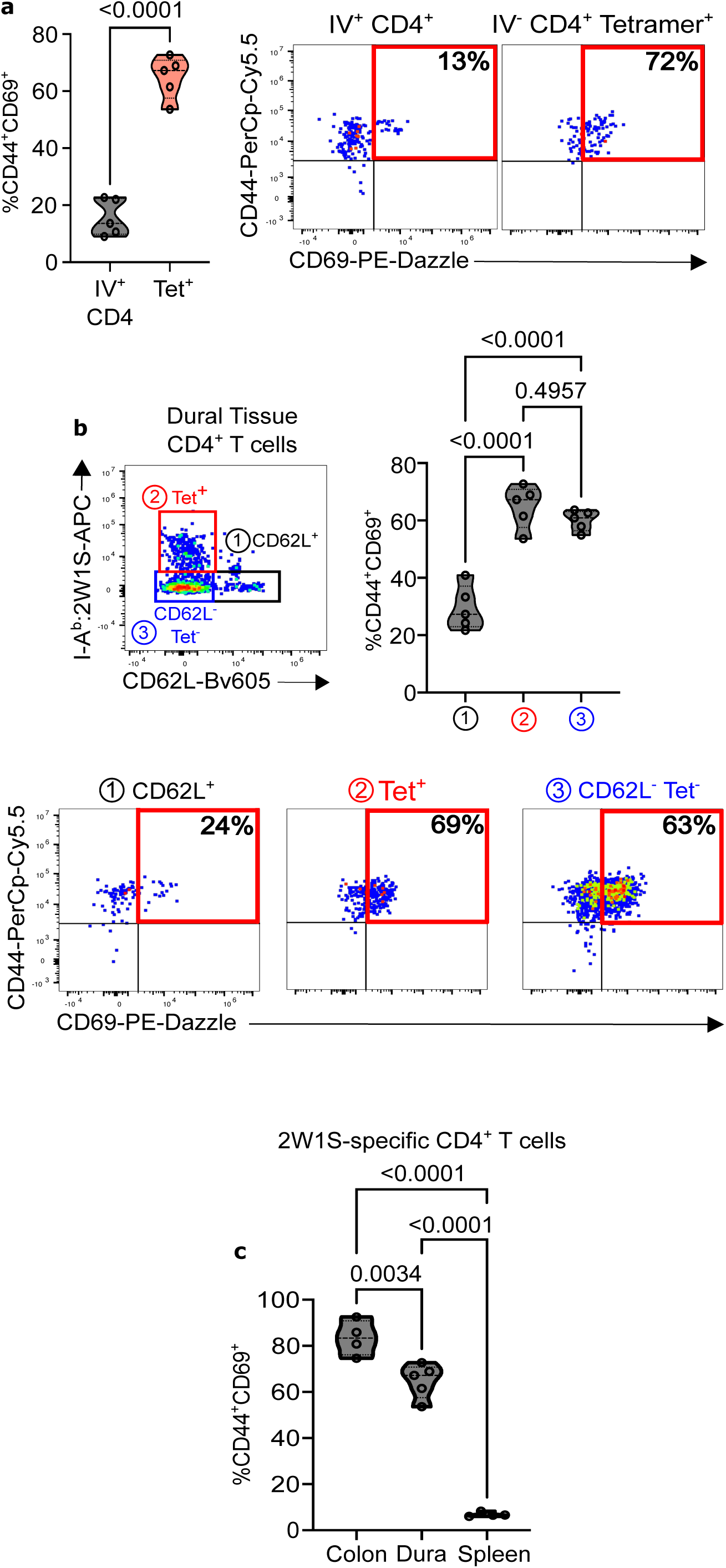
(related to Main Figure 3 D-E) **A:** Cumulative violin plots (left) and representative cytometry (right) of CD44/CD69 expression on dural intravascular CD4^+^ T cells (“IV+ CD4+”) *versus* I-A^b^:2W1S tetramer-binding T cells (“IV-CD4^+^ tetramer^+^) from dural tissue following oral *S.* typhimurium infection (21 dpi). *P* value calculated by student’s unpaired *t* test **B**: Cumulative violin plots (top) and representative cytometry (bottom) showing expression of CD44/CD62L on the indicated cell subsets following oral *S.* typhimurium infection (1 = naïve T cells CD62L^-^, 2 = 2W1S antigen-specific, 3 = non-2W1S-specific antigen-experienced T cells). **C**: Representation of CD44^+^CD69^+^ cells amongst total 2W1S antigen-specific CD4^+^ T cells from the indicated organs following oral *S.* typhimurium infection. *P* values in **B** and **C** calculated using ordinary one-way ANOVA with Tukey’s multiple comparisons test

**Supplemental Figure 5.**
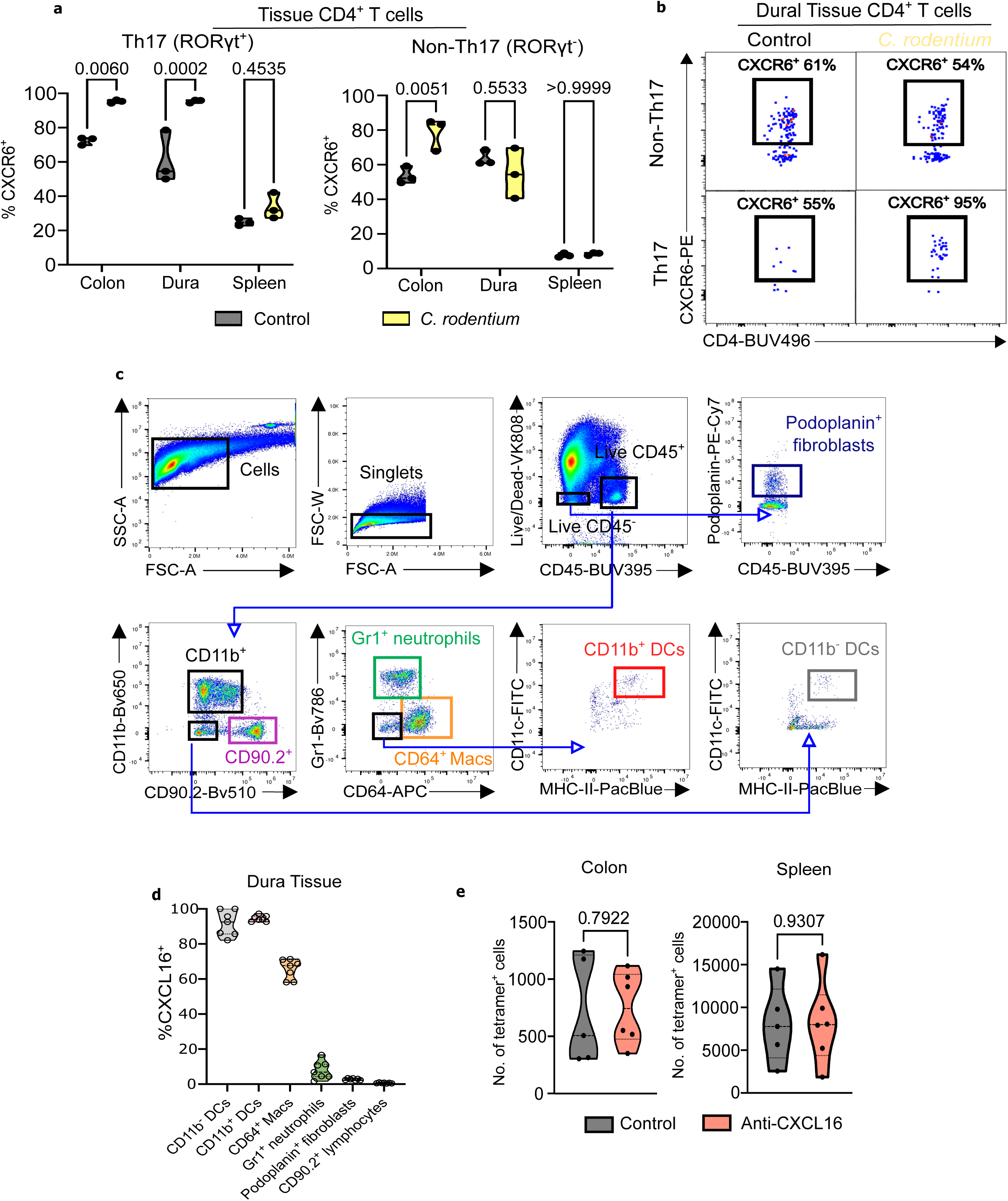
(related to Main Figure 4 B, E-H) **A:** Representation of CXCR6^+^ cells amongst total Th17 (left) and non-Th17 CD4^+^ (right) T cells from the specified organs (*n* = 3). two-way ANOVA with Bonferroni’s post-test **B**: Representative cytometry showing CXCR6 expression amongst dural Th17 and non-Th17 CD4^+^ T cells in *C. rodentium-*infected *versus* control mice. (21 dpi). **C**: Gating strategy to identifying dural myeloid, lymphoid, and fibroblast populations for CXCL16 staining. **D**: Representation of CXCL16^+^ cells amongst the indicated cell subsets from dural tissue *ex vivo* (*n* = 7). **E**: Bead-normalised quantification of I-A^b^:2w1s tetramer-binding T cells from the colon and spleen of *S.* typhimurium-2W1S-infected mice receiving anti-CXCL16 *versus* control (*n* = 5-6 per group). *P* values derived by unpaired student’s *t*-test

**Supplemental Figure 6.**
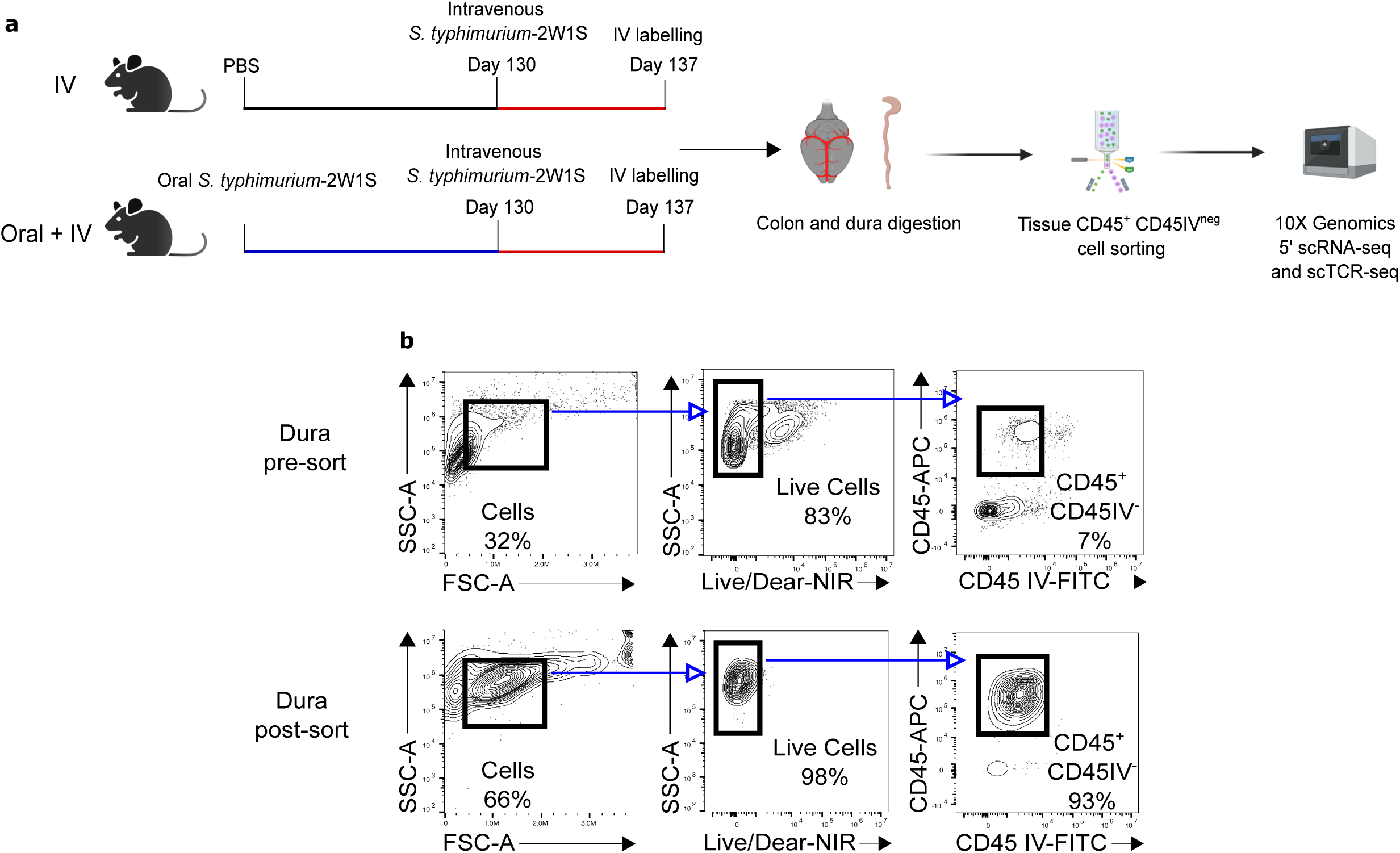
(related to Main Figure 5 E) **A:** Schematic of experimental setup for primary (oral) and secondary (IV) *S.* typhimurium infection for single-cell RNA sequencing and single-cell TCR sequencing. **B**: Representative cytometry of dural immune cells before (top) and after (bottom) cell sorting for scRNA-seq experiment.

**Supplemental Figure 7.**
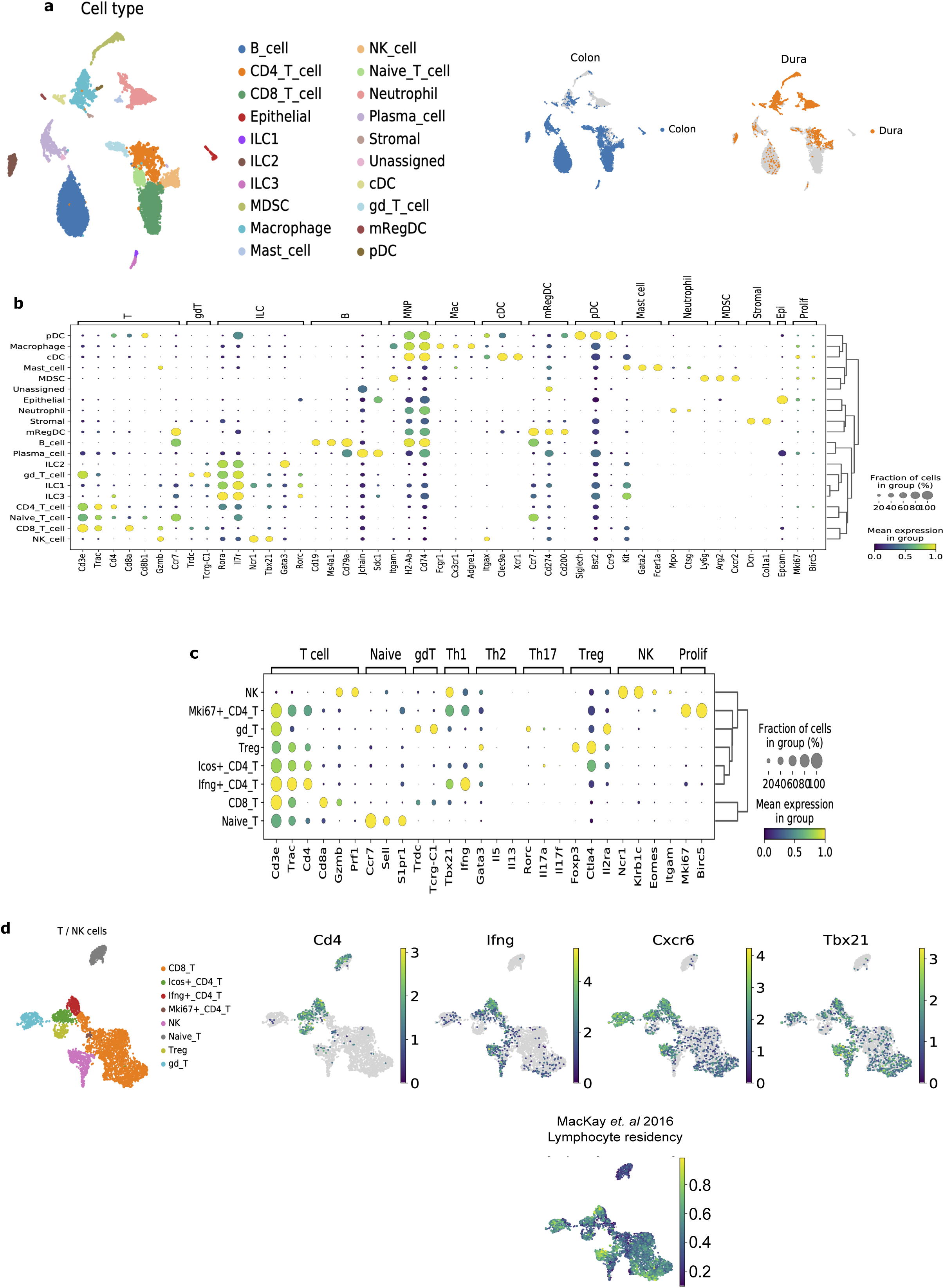
(related to Main Figure 5 E-H) **A**: UMAP of 11655 cells obtained from scRNA-seq of extravascular CD45+ immune cells in the dura or colon, coloured by cell type and tissue origin **B**: Dot plot of canonical marker gene expression of cell types in (A) **C**: Dot plot of canonical marker genes and selected differentially expressed genes of cell types in Fig. 5 (d) **D**: (left) UMAP of T and NK cells (subset from (a)) showing T and NK cell subsets and (right) UMAP of selected expressed CD4^+^ Th1 genes and lymphocyte residency markers from MacKay *et. al* (2016)

**Supplemental Figure 8.**
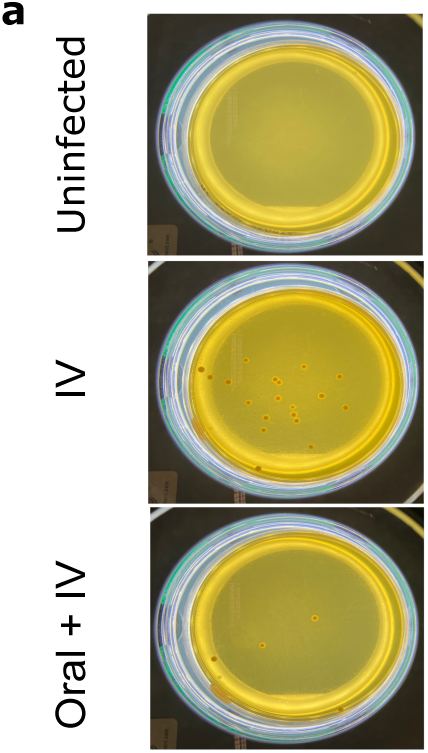
(related to Main Figure 5 K) **A:** Representative YPD plates of *C. albicans* growth in the different conditions as specified in Fig. 5 (k).

